# Genetic Evidence for Geographic Structure within the Neanderthal Population

**DOI:** 10.1101/2023.07.28.551046

**Authors:** Alan R. Rogers

**Affiliations:** University of Utah

## Abstract

PSMC estimates of Neanderthal effective population size (*N*_*e*_) exhibit a roughly 5-fold decline across the most recent 20 ky before the death of each fossil. To explain this pattern, this article develops new theory relating genetic variation to geographic population structure and local extinction. It argues that the observed pattern results from subdivision and gene flow. If two haploid genomes are sampled from the same subpopulation, their recent ancestors are likely to be geographic neighbors and therefore coalesce rapidly. By contrast, remote ancestors are likely to be far apart, and their coalescent rate is lower. Consequently, *N*_*e*_ is larger in the distant past than in the recent past.

New theoretical results show that modest rates of extinction cause substantial reductions in heterozygosity, Wright’s *F*_*ST*_, and *N*_*e*_.

## 1 Introduction

A variety of statistical methods use genetic data to estimate the history of effective population size, *N*_*e*_ [12, 21, 22, 24, 32]. In a subdivided population, these estimates depend not only on the number of individuals but also on gene flow between subdivisions [19, 34, 38]. Not only does *N*_*e*_ change in response to changes in gene flow [17, 20, 33], it may also exhibit a prolonged decline even when there has been no change either in the number of individuals or in the rate or pattern of gene flow [18, 20].

With this in mind, consider the data in Fig. 1, which replots previously-published estimates of archaic population histories [13]. This figure zooms in on an interval of 30 ky before the death of each fossil. PSMC estimates are famously unreliable over this time scale [12]. It is therefore unsurprising that the Denisovan curve swings wildly up and down, a pattern consistent with statistical error. On the other hand, the three Neanderthal curves seem to tell a consistent story—one of a roughly 5-fold decline in population size across 20 thousand years. The consistency of the Neanderthal curves is surprising and demands explanation.

**Figure 1:**
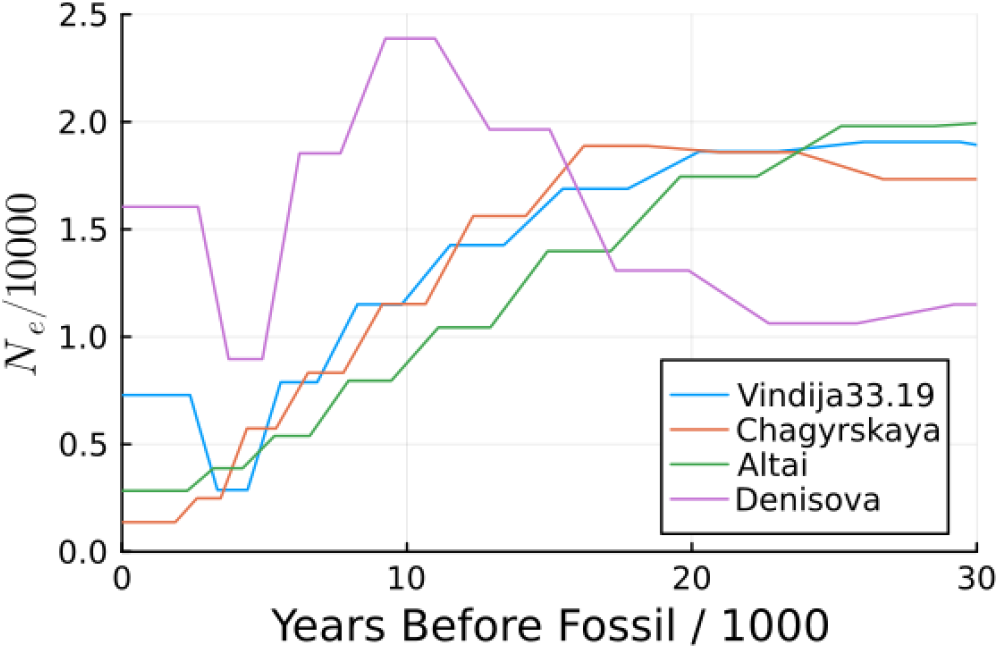
PSMC estimates of the recent history of population size based on three Neanderthals (Vindija33.19, Chagryskaya, and Altai) and a Denisovan [13]. Horizontal axis measures time in thousands of years before the date of each fossil. Mutation rate is 1.4 × 10^*−*8^ per base pair per generation; generation time is 29 y.

In what follows, I ask whether this pattern may reflect geographic subdivision within the Neanderthal population rather than either statistical noise or a real decline in the number of Neanderthals.

## 2 Methods

### 2.1 Archaic PSMC data

The data in Fig. 1 were published by Mafessoni et al. [13] and were provided by those authors. I used psmcdata to extract data from PSMC output files.

### 2.2. Coalescent hazard and effective population size

The effective population size, *N*_*e*_(*t*), at time *t* is defined in terms of the corresponding hazard, *h*(*t*), of a coalescent event. To understand this latter quantity, suppose that we sample two genes from a population at time 0 and trace the ancestry of each backwards in time. Eventually, the two lineages will *coalesce* into a common ancestor [6, 31]. The *hazard, h*(*t*), of a coalescent event at time *t* is the conditional coalescent rate at *t* given that the two lineages did not coalesce between 0 and *t* [8, p. 6]. In a randomly-mating population of constant size *N*, this hazard is *h* = 1*/*2*N* per generation. In a population of more complex structure, this formula doesn’t hold. Nonetheless, we can define an *effective population size* as half the reciprocal of the hazard ([11]; [23, p. 144]).

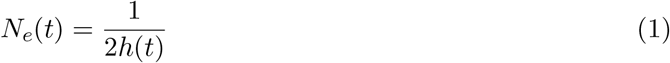

This quantity has also been called the inverse instantaneous coalescent rate (IICR) [4, 17, 18, 20]. That alternative name emphasizes that change in *N*_*e*_ (or IICR) need not imply change in the size of the population or even in the rate or pattern of mobility within it. Furthermore, this quantity depends not only on the characteristics of the population but also on those of the sample. Thus, the conventional term—effective population size—may be misleading. Nonetheless, I adhere to that convention in this article.

Because so little is known about subdivision and gene flow within the Neanderthal population, we cannot hope to build a realistic model. Instead, I use simple models to explore the effect of subdivision, gene flow, and local extinction (or extirpation). Extinction is important because although the Neanderthal population was widespread and shows evidence of geographic structure, it had low heterozygosity [13]. If demes never went extinct, population structure would tend to inflate heterozygosity ([38, p. 133]; [19]). But the opposite happens when demes occasionally go extinct and are then replaced by immigrants from another deme ([37, p. 244]; [26]; [14]; [35]; [34]). Thus, the low heterozygosity of Neanderthals suggests that local extinctions may have been common.

### 2.3 The finite island model with local extinction

Consider first the finite island model, which assumes that demes of equal size exchange migrants at equal rates [3, 17, 20]. In reality, a gene beginning in one deme may need to traverse many intervening demes to reach one that is far away. There is no such necessity under the island model: a gene can move from one deme to any other in a single generation. Consequently, this model converges rapidly toward its asymptote for any given level of gene flow.

The finite island model assumes *d* demes of effective size *N*. Pairs of lineages within the same deme coalesce at rate 1*/*2*N* ; pairs in different demes cannot coalesce. Migration occurs at rate *m* per gene per generation. When a migration occurs, the gene moves to one of the *d* − 1 other demes. In addition to these standard assumptions, I also assume that demes occasionally go extinct and are then immediately replaced with immigrants from some other deme. In backwards time, extinctions look like wholesale migration: all lineages within the deme migrate together to a different deme, which is chosen at random. Extinctions occur at rate *x* per deme per generation.

As we trace the history of a pair of genes, we need only keep track of whether the two lineages are colocal (in the same deme) or in different demes. Extinction can be ignored for a pair of colocal lineages. It is not that extinctions don’t happen. As we view a pair of colocal lineages at the time of an extinction, they move together to another deme, but in that new deme they are still colocal. The state of the system is thus unchanged. On the other hand, extinction does affect pairs in different demes. At the time of the extinction, one lineage moves to a different deme, and this new deme may be the one occupied by the other lineage. Thus, extinction increases the rate at which separated lineages become neighbors.

Other models of local extinction [e.g. 26] have included temporary reductions in population size (bottlenecks) at the time of recolonization, which reduce the effective size of each deme. Rather than modelling these explicitly, I absorb them into effective deme size, *N*. If bottlenecks are frequent, *N* will be smaller than the average size of a deme.

To study this model, let us formulate it as a Markov chain in continuous time. At a given time in the past, the two sampled lineages will be in one of three states: *S*, they are in the same deme; *D*, they are in different demes; and *C*, they have coalesced into a single lineage. Once the process reaches state *C*, it never leaves, so the probability of this state increases through time. Such states are said to be *absorbing*. States *S* and *D*, on the other hand, are *transient* ; their probabilities will eventually decline toward zero. Let *q*_*ij*_ denote the rate of transitions from state *i* to state *j*. In other words, *q*_*ij*_ *δt* is approximately the probability that the process is in state *j* at time *t* + *δt* given that it is in state *i* at time *t*, provided that *δt* is small. The matrix of transition rates looks like

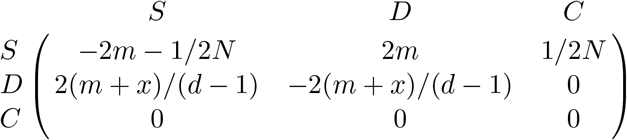

Here, 2*m* is the rate of transition from *S* to *D* (because there are two lineages, each of which migrates at rate *m*), and 1*/*2*N* is the rate of transition from *S* to *C*. In other words, it’s the rate of coalescence for two lineages in the same deme. Note that *x* (the rate of extinction) does not contribute to transitions from state *S*. On the other hand, extinction and migration both contribute to the rate of transition from *D* to *S*. These transitions occur at rate 2(*m* + *x*)*/*(*d* − 1), because although 2(*m* + *x*) is the combined rate of migrations and extinctions, only a fraction 1*/*(*d* − 1) of these events results in one lineage joining the other in the same deme. The diagonal entries in a transition rate matrix are the negative of the sum of the other entries in that row. (See Sukhov and Kelbert [30s, sec. 2.1] for a discussion of transition rate matrices.) If *x* = 0, this matrix is equivalent to that of Rodríguez et al. [20, p. 668].

We can simplify this model in two ways. First, because the absorbing state, *C*, does not contribute to the probabilities of the other states, we can delete the third row and column to form a *subintensity matrix* [2, p. 125]. Second, we can reexpress all rates using 2*N* generations as the unit of time. This involves multiplying all entries of the matrix above by 2*N*. The result is

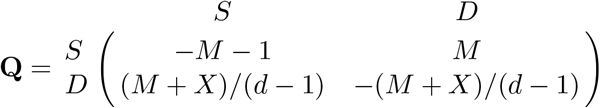

where *M* = 4*Nm* is the migration rate per pair of lineages per 2*N* generations, and *X* = 4*Nx* is the corresponding rate of extinction. Let *τ* = *t/*2*N* represent time in units of 2*N* generations, and let **p**(*τ*) = (*p*_1_(*τ*), *p*_2_(*τ*)) represent the row vector of probabilities that the process is in states *S* and *D* at time *τ*. It equals

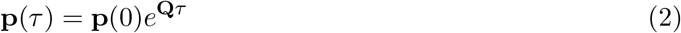

where **p**(0) is the vector of initial probabilities, and *e*^**Q***τ*^ is a matrix exponential [5s, p. 182]. Because we have sampled two genes from the same deme, **p**(0) = (1, 0), and **p**(*τ*) is the first row of *e*^**Q***τ*^.

As *τ* increases, both entries of **p**(*τ*) will eventually decline toward zero, because it becomes increasingly unlikely that the two lineages have not yet coalesced. We are interested, however, in the conditional probability that the two are in the same deme, given that they have not yet coalesced. At time *τ*, this conditional probability is

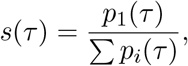

where the sum is across the transient states, excluding state *C*. The coalescent hazard is the product of *s*(*τ*) and the hazard for two lineages in the same deme. Returning now to a time unit of one generation, that product is *h*(*t*) = *s*(*t/*2*N*) */*2*N*, and effective population size is

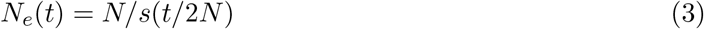

This differs from previously published results on the IICR ([17, Eqn. 16]; [20, p. 669]) only in that it adds the effect of local extinctions.

### 2.4 The circular stepping stone model with local extinction

One-dimensional stepping stone models describe a population whose demes are arranged in a line and exchange genes only with their immediate neighbors [10]. They are abstractions of real-world situations in which demes are arrayed across a landscape, and neighboring pairs of demes are less isolated from each other than are pairs separated by large distances. I will follow the common practice of assuming that the ends of the line of demes are joined to form a circle of *d* demes, each of effective size *N*, and that each pair of neighboring demes exchanges migrants at the same rate [3, 15, 16]. When local extinctions occur, the lost deme is immediately replaced with immigrants from a single donor deme. The donor deme is equally likely to be either of the two adjacent demes. The circular arrangement is for convenience only. It simplifies things, because it implies that no deme is more central or peripheral than any other.

Each lineage migrates at rate *m* per generation in backwards time, and when it does so it is equally likely to move one step clockwise or one step counterclockwise around the circle of demes. Because we are studying the history of a pair of genes, the migration rate is 2*m* per generation or *M* = 4*Nm* per unit of 2*N* generations, provided that the two lineages have not yet coalesced. Similarly, if two lineages are in different demes, 2*x* is the rate per generation at which extinction affects one of them or the other, and *X* = 4*Nx* is the rate per 2*N* generations.

As in the island model, this process has one absorbing state, *C*, in which the ancestors of the two sampled genes have coalesced. Two lineages are separated by 0 steps if they are in the same deme, by 1 step if they’re in adjacent demes, and so on. The maximum separation is ⌊*d/*2⌋, the largest integer less than or equal to *d/*2. For example, ⌊*d/*2⌋ = 3 if *d* is either 6 or 7. The transient states in the model correspond to these distances: 0, 1, …, ⌊*d/*2⌋. These states label the rows and columns of the subintensity matrices below. The matrix is

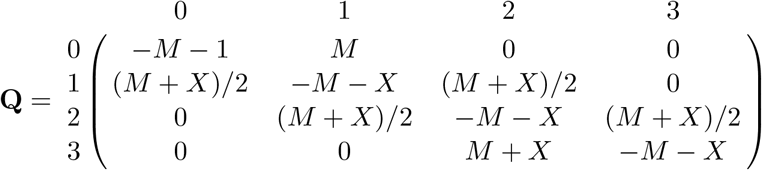

for *d* = 6 and

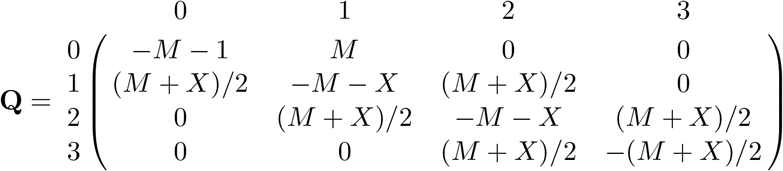

for *d* = 7. In each matrix, row 0 refers to the case in which the two lineages are in the same deme. The only positive entry in that row equals *M*, because extinction can be ignorned, and any migration in state 0 will move the process to state 1. The “–1” in the left-most entry accounts for coalescent events. In rows 1 and 2, the two positive entries equal (*M* + *X*)*/*2, because in states 1 and 2, migration and extinction are equally likely to increase by 1 or to reduce by 1 the distance between lineages. The entries in row 3 depend on whether *d* is even or odd. If it is even, there is only one deme that is ⌊*d/*2⌋ steps away from any given deme. Consequently, any migration or extinction must reduce the distance by 1 step, and the transition rate is *M* + *X*. On the other hand, if *d* is odd, there are two demes ⌊*d/*2⌋ steps away. Half of migration and extinction events will move a lineage from one of these to the other without changing the distance between demes. The other half reduce that distance by 1, so the transition rate is (*M* + *X*)*/*2. *N*_*e*_(*t*) is calculated from **Q** as before, using Eqns. 2–3.

### 2.5 Genetic variation within and among groups

The “within-group heterozygosity,” *H*_*W*_, is the probability that two genes drawn at random from the same deme differ in state. The appendix shows that

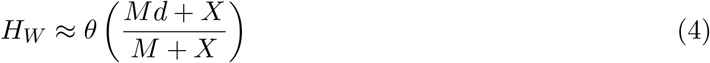

under either of the two models described above. If *X* is similar in magnitude to *M* or is larger, extinction produces a substantial reduction in heterozygosity. When *X* = 0, this reduces to a well-known result for models without extinction [27, 29].

The appendix also derives formulas for Wright’s *F*_*ST*_ under each model. For the island model,

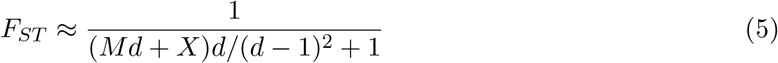

When *X* = 0, this reduces to a well-known formula for *F*_*ST*_ under the island model [28]. For the circular stepping-stone model,

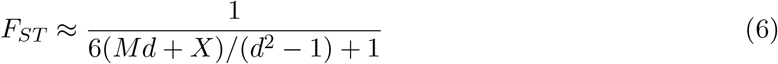

When *X* = 0, this result is equivalent to that of Wilkinson-Herbots [36, p. 582], whose results suggest a caveat. She presents two results. One (which Eqn. 6 generalizes) is an approximation for weak mutation [36, p. 582]. The other [36, Eqn. 34] is a more accurate formula that includes a mutation rate. With a realistic mutation rate and a modest number demes, her two formulas give nearly identical results. But as the number, *d*, of demes increases, the two formulas diverge. For large *d*, the process apparently generates such long coalescent times that the weak-mutation approximation breaks down. This is probably also true of my Eqn. 6.

### 2.6 The magnitude of change in *N*_*e*_(*t*)

For both of the models discussed above, *N*_*e*_(*t*) increases toward an asymptote in backwards time. The asymptotic value, *N*_*e*_(∞), can be obtained from the left eigenvector of **Q** associated with the largest eigenvalue. This eigenvector is proportional to the asymptotic value of **p**(*τ*), so we can use it to calculate *N*_*e*_(∞), just as we used **p**(*τ*) to calculate *N*_*e*_(*t*). If 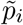 is the *i*’th entry of this eigenvector, then the asymptotic value is 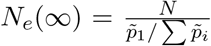, where 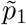 is the first entry of the eigenvector and corresponds to state in which both lineages are in the same deme. Mazet et al. [17] derived an explicit formula for the asymptote under the island model. The maximum proportional increase as we move backwards in time is

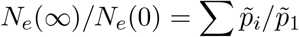

### 2.7 Computer simulations

Computer simulations were done using Msprime [1, 9]. This software has no support for random extinctions, so my program runs its own simulation to generate a list of extinctions and recolonizations, which is then used to build the demographic model of Msprime. Each run of Msprime uses a different, randomly-generated list of extinctions and recolonizations.

## 3 Results

This section asks whether geographic population structure can account for the Neanderthal pattern in Fig. 1. To this end, it fits each of the models described above to four observations. The first two of these come from Fig. 1: (1) a decline in *N*_*e*_ over a period of about 20 ky, ending at the time of the Neanderthal fossil; and (2) the ratio of early to late *N*_*e*_ is roughly 5 or 6. In addition: (3) Neanderthals had very low heterozygosity, as discussed below; and (4) *F*_*ST*_ in vertebrate species is usually less than 0.5 [25]. Having shown that all of these observations can be explained by population structure, I then ask whether the pattern in Fig. 1 can be explained as an artifact of sampling or as a real decline in the number of Neanderthals.

To constrain the models, let us begin with heterozygosity. Mafessoni et al. [13, table S8.2] list values between 1.5 × 10^*−*4^ and 2 × 10^*−*4^ for four archaic genomes. I take the upper end of this range as representative of Neanderthal heterozygosity. I also assume a mutation rate of 1.4 × 10^*−*8^ per nucleotide site per generation. If local demes never went extinct (i.e., if *X* = 0), we could plug these assumptions into Eqn. 4 and find that the sum of effective sizes of Neanderthal subpopulations was only about 3600. This value seems implausibly low for a population as widespread as Neanderthals, so I will assume that *X* equals the migration rate, *M*. This value is large enough to roughly double the implied size of the metapopulation. Nonetheless, it is still a modest rate of extinction, as discussed below. With these assumptions, one can set *H*_*W*_ equal to observed heterozygosity and solve for *d*, the number of demes, as a function of deme size, *N*.

For any choice of *N* and *d*, we can make graphs like those in Fig. 2. These graphs (and many others not shown) all exhibit declines in *N*_*e*_, which do not reflect any real decline in the number of individuals. Instead, they are the effect of sampling two haploid genomes from a single deme within a structured population. The rate of decline depends on *M*, and within each graph we will be interested in the value of *M* that most closely matches the 20-ky decline seen in Neanderthal data (Fig. 1). This suggests *M* = 6 for the island model and *M* = 20 for the stepping-stone model. These choices generate declines in *N*_*e*_ over the right interval. These declines also have roughly the right magnitude: *N*_*e*_(∞)*/N*_*e*_(0) ≈ 7 for the island model and 6 for the circular stepping-stone model. Finally, they imply reasonable *F*_*ST*_ values: 0.11 for the island model and 0.08 for the circular stepping-stone model.

**Figure 2:**
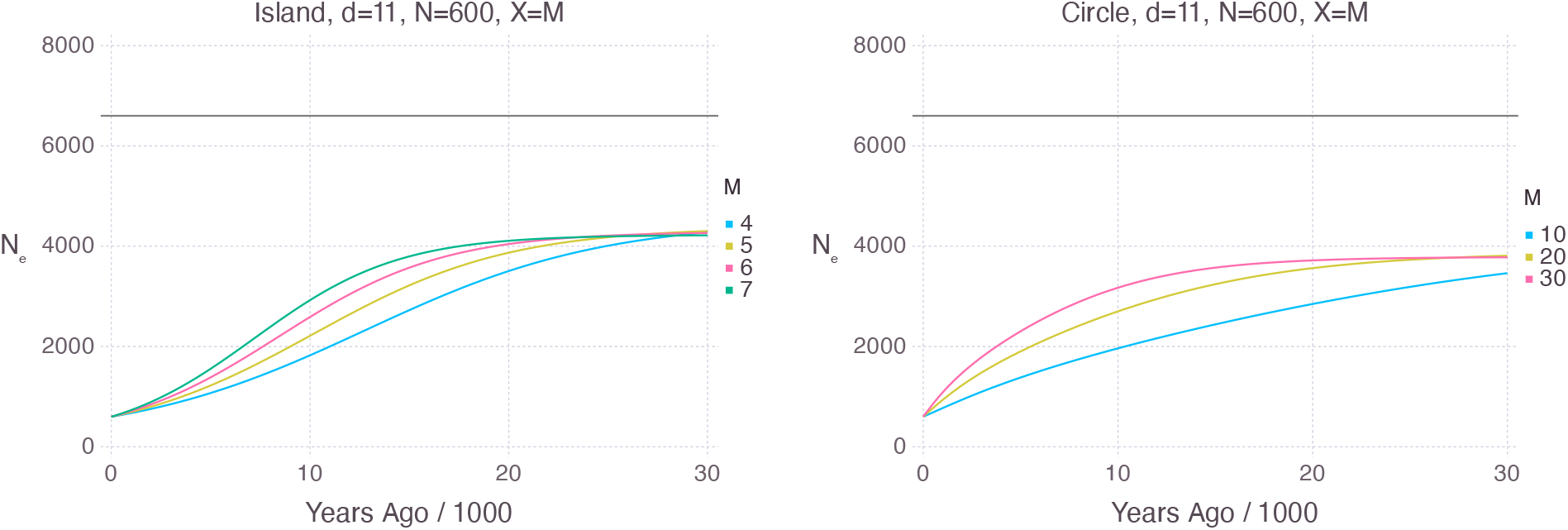
Time path of *N*_*e*_ in models with extinction, assuming that two genes are sampled from the same deme and that the generation time is 29 y. The horizontal gray line shows the metapopulation size, *Nd*.

Let us re-express the inferred rates of migration and extinction in terms that are easier to interpret. For the island model, *M* = 6 means that each deme receives *M/*4 = 1.5 immigrants per generation. Because *N* = 600 in these models, *X* = 6 means that the extinction rate is *X/*4*N* = 1*/*400 per deme per generation and that the mean interval between extinction events is 400 generations. For the circular stepping-stone model, the corresponding values are 5 immigrants per deme per generation and 120 generations between extinction events. These are modest rates of extinction, and the true rate may have been larger. If so, the Neanderthal metapopulation would have been larger than the value implied here. It would have been larger still if local groups experienced frequent bottlenecks, perhaps associated with recolonization events. Such bottlenecks would make effective deme size, *N*, smaller than census size. Consequently, the census size of the metapopulation would be larger than *Nd*.

There are many other choices of parameter values that also fit the data. For example, Fig. 3 looks at two versions of the island model, one with no extinction (*X* = 0) and the other with a high rate (*X* = 10*M*). Both fit the data best if we assumue that *M* = 6, but the two models have drastically different implications for the total size, *Nd*, of the metapopulation: that size is 3600 if we assume no extinction but 33,000 if we assume a high rate of extinction. Even with an extinction rate this high, the interval between extinction events is still 40 generations—slightly more than 1000 y—and does not seem implausible. Thus, the low Neanderthal heterozygosity does not imply that their global population was small.

**Figure 3:**
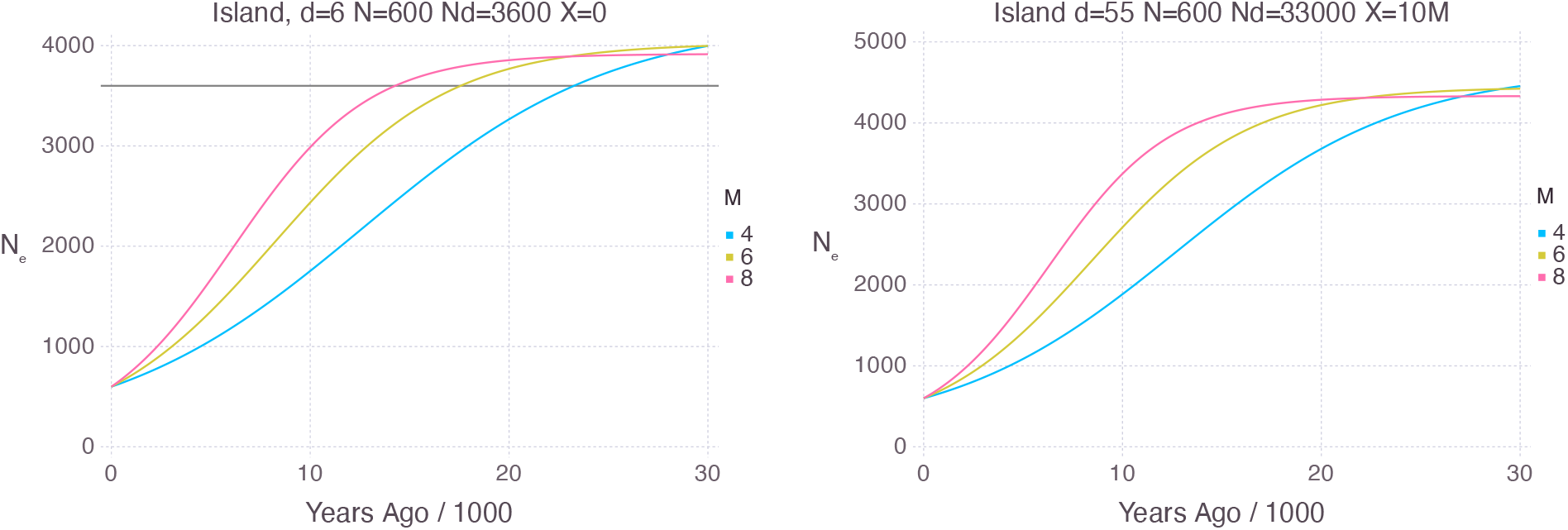
Island model with low (*X* = 0) and high (*X* = 10*M*) rates of extinction. For *M* = 6, *F*_*ST*_ is 0.10 on the left and 0.12 on the right. *N*_*e*_(∞)*/N*_*e*_(0) is 6.7 on the left and 7.4 on the right. The sum, *Nd*, of effective deme sizes is 3600 on the left and 33,000 on the right. The horizontal line in the left panel is the sum, *Nd*, of effective deme sizes.

There is nothing special about the island and circular stepping-stone models. These represent two extremes—one in which isolation by distance is absent and another in which it takes an extreme form. Because both extremes are consistent with the data, it seems likely that intermediate models would also work. The present results tell us only that the Neanderthal data are consistent with geographic population structure.

What other mechanisms might explain the data? We should worry first about statistical error. PSMC is notoriously unreliable [12] over the short time scale graphed in Fig. 1. Perhaps the data are a statistical fluke. This hypothesis is hard to reconcile with the consistency of the three Neanderthal curves. On the other hand, consistency is to be expected under the hypothesis of population structure. Each panel of Fig. 4 shows three simulated replicates, which are quite similar to each other, and also to the curves in Fig. 1.

**Figure 4:**
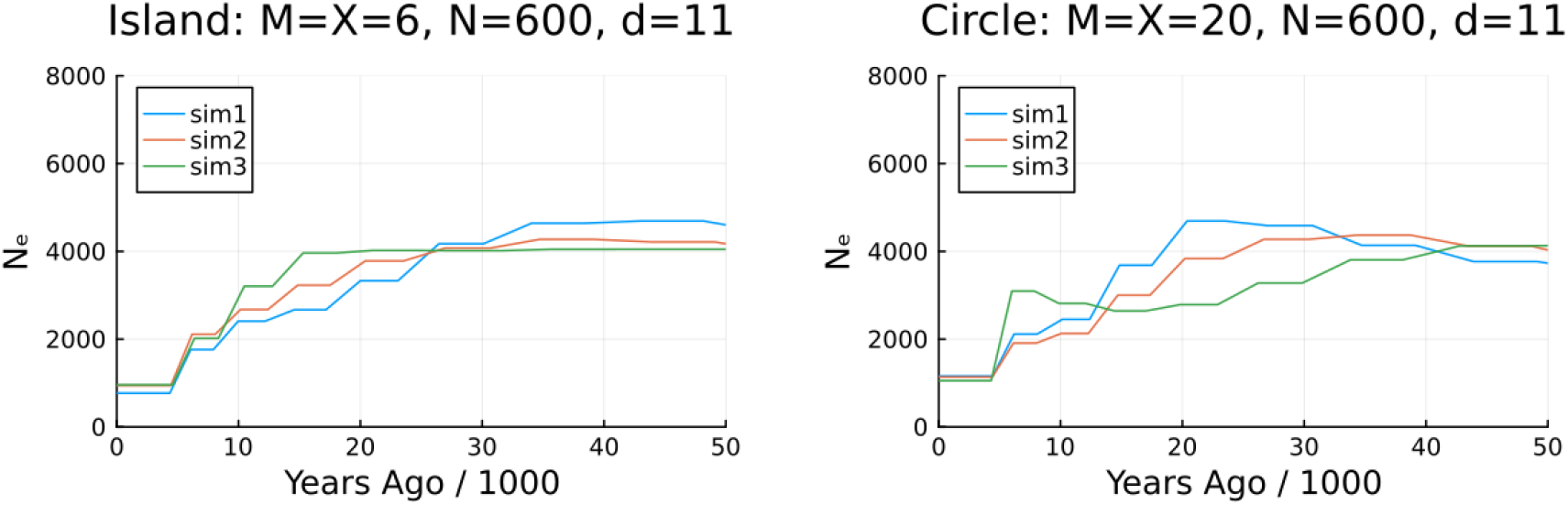
PSMC estimates of *N*_*e*_ based on simulated data. Simulations used Msprime [1, 9] and assumed 11 demes of size 600. Genomes comprised 10 pairs of chromosomes, each 10^7^ base pairs in length. Rates of migration and extinction (*M* = *X* = 6 for island model and *M* = *X* = 20 for stepping stone) were chosen using Fig. 2 to produce a decline in *N*_*e*_ over about 20 ky.

On the other hand, perhaps the pattern in Fig. 1 reflects a real decline in Neanderthal population size. The problem with this hypothesis is that the three Neanderthal fossils lived at different times. The youngest (Vindija) lived 60 ky after the oldest (Altai) [13, p. 1532]. If these PSMC curves were recording the same decline in the size of the Neanderthal metapopulation, the declines should not appear simultaneous in Fig. 1; they should be separated by 60 ky. This is obviously not the case, so it seems unlikely that the observed pattern reflects a real decline in the Neanderthal metapopulation. The synchrony of these curves is not a problem under the hypothesis of population structure. That hypothesis implies that *N*_*e*_ will increase gradually in backwards time, beginning at the date of each fossil. The similar rates of increase suggest that similar rates of gene flow prevailed for all three Neanderthal fossils.

## 4 Discussion

Other authors [4, 17, 18, 20] have emphasized that a decline in effective population size, *N*_*e*_, may result from geographic population structure, even without any change in census population size or in the rate or pattern of gene flow. Figs. 2 and 3 show how this works in the context of two theoretical models. The declines in *N*_*e*_ reflect the fact that we have sampled two genes from a single deme. In the recent past, it is likely that the two ancestors of these genes are still neighbors. Coalescent hazard is therefore high and *N*_*e*_ is small. As we move farther into the past, ancestors are less likely to be neighbors, coalescent hazard declines, and *N*_*e*_ increases toward an asymptote.

The position of this asymptote depends on rates of migration and extinction. If demes never go extinct, the asymptote, *N*_*e*_(∞), is even larger than *Nd*, the size of the metapopulation. (See the horizontal gray line in the left panel of Fig. 3.) Why should the asymptote be so large? Among all possible histories of a pair of genes, the subset that has not yet coalesced at time *t* will be enriched with histories in which the two lineages haven’t spent much time together in the same deme. This implies that when *t* is large, the two lineages are less likely to be in the same deme than are two genes drawn at random from the population as a whole. Consequently, coalescent hazard is less than 1*/*2*Nd*, and *N*_*e*_(∞) *> Nd*. This excess is pronounced if the migration rate, *M*, is small but disappears as *M* grows large. On the other hand, if demes do occasionally go extinct, the asymptote is smaller, and *N*_*e*_(∞) may be smaller that *Nd*, as seen in Fig. 2. In this figure, the extinction rates are such that demes persist for hundreds of generations. In the right panel of Fig. 3, the extinction rate is higher so that demes persist only for 40 generations. In such cases, the metapopulation is dramatically larger than the *N*_*e*_ that we estimate.

It seems likely that population structure underlies the apparent decline in Neanderthal *N*_*e*_ shown in Fig. 1. Alternative explanations are unable to account for the consistency of the three Neanderthal curves or for the fact that they all begin at the same point on the horizontal axis. This conclusion supports that of Mafessoni et al. [13], who use runs of homozygosity to argue that the Neanderthal population was geographically structured.

Although the Denisovan curve in Fig. 1 does not exhibit the decline characteristic of population structure, I would not argue that this population lacked structure. In spite of the consistency of the curves in Fig. 4, such simulations do generate aberrant curves reasonably often. We should not read too much into a single empirical curve. Furthermore, there is convincing evidence that the Denisovan population was structured [7].

## 5 Conclusions

PSMC estimates of Neanderthal population size exhibit a consistent roughly 5-fold decline during the most recent 20 kyr. The consistency of these estimates is surprising, both because PSMC is thought to be unreliable on this time scale and also because we are measuring time backwards from three fossils that lived at very different times. The observed pattern does not seem to be an artifact of sampling, nor is it likely to reflect a real decline in the size of the Neanderthal metapopulation. Instead, it is supports a hypothesis of geographic population structure.

This article presents mathematical theory describing the effect of geographic population structure and local extinctions on *N*_*e*_, *F*_*ST*_, and heterozygosity. This theory shows that if extinction rates were such that local Neanderthal populations persisted for only 1000 y or so, the global Neanderthal population would have been dramatically larger than its effective size.

## Acknowledgements

I am grateful to Fabrizio Mafessoni for providing the data in Fig. 1 and for comments from Simon Boitard, Elizabeth Cashdan, Lounès Chikhi, Josué Corujo, Olivier Mazet, and Daniel Tabin. Jerome Kelleher provided helpful advice on Msprime.

## Data, script, and code availability

Source code is available at doi:10.17605/OSF.IO/T4N3C. The latest version of this code is available at https://github.com/alanrogers/StrucPopMod.

## A Two models of geographic population structure

This section studies two models of geographic population structure: the finite island model and the circular stepping stone model. Both assume a metapopulation of *d* demes, each of effective size *N*. Time is measured in units of 2*N* generations. On this time scale, pairs of lineages in the same deme coalesce at rate 1. If each lineage mutates at rate *u* per generation, then 2*Nu* is the rate per 2*N* generations. When we trace the history of a pair of lineages in continuous time, we can ignore the possibility that both lineages mutate in the same instant. When mutation occurs, one lineage or the other is affected, and this happens at rate *θ* = 4*Nu*—twice the rate of an individual lineage. Similarly, if *m* is the rate of migration per lineage per generation, then *M* = 4*Nm* is the rate per 2*N* generations for a pair of lineages. If *x* is the extinction rate per deme per generation, then *X* = 4*Nx* is the rate per 2*N* generations for a pair of lineages in separate demes.

To explore the two models below, I use Slatkin’s approximation [28], which expresses heterozygosity or gene diversity in terms of mean coalescence time. This approximation is based on the following idea. If *f* (*t*) is the probability density that two genes coalesce at time *t*, then they are identical in state with probability

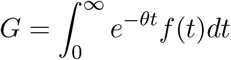

where *t* is time in units of 2*N* generations, and *e*^*−θt*^ is the probability that neither lineage mutates in the interval between 0 and *t*. If *θ* is small, then *e*^*−θt*^ ≈ 1 − *θt*, and 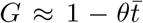, where 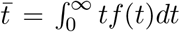 is the mean coalescence time. Under this approximation, heterozygosity within demes— the probability that two genes sampled from the same deme differ in state—is

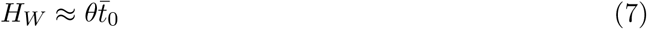

where 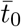 is the mean coalescence time for pairs of genes sampled within a single deme. Wright’s *F*_*ST*_, a measure of differentiation among demes, is [28, p. 169]

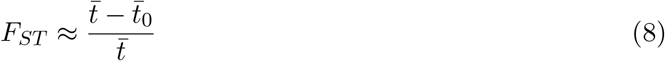

where 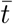 is the mean coalescence time for two genes drawn from the population as a whole.

A.1 The finite island model with extinction

In this section, 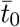 represents the expected coalescence time, in units of 2*N* generations, for a pair of genes sampled from the same deme. For pairs sampled from different demes, the analogous quantity is 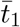 Pairs in the same deme coalesce at rate 1 and migrate at rate *M* per unit of 2*N* generations. Extinction can be ignored for pairs in the same deme. Thus,

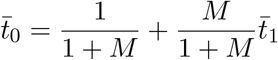

The first term on the right is the expected time until an event of either type. If that event is a coalescence, then we are done: there are no further contributions to 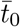. But with probability *M/*(1+*M*), the event is a migration, the two lineages are now in separate demes, and their expected coalescence time becomes 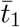.

Pairs in different demes cannot coalesce but are affected by migration (rate *M*) and extinction (rate *X*). When either of these events occur, the two lineages end up in the same deme with probability 1*/*(*d* − 1). Thus, two separated lineages join each other in the same deme at rate (*M* + *X*)*/*(*d* − 1) and

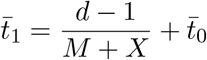

Here, the first term is the expected time until a pair in different demes moves into the same deme. Substitute the formula for 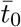 into this expression and solve for 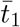. The result is

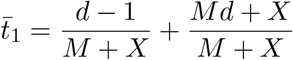

Comparison of the two expressions for 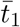 shows that 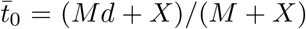. Substituting this into (7) gives Eqn. 4. The mean coalescence time of two genes drawn at random from the entire metapopulation is

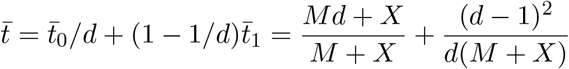

Substituting into (8) gives Eqn. 5.

### A.2 Circular stepping-stone model with extinction

If two lineages are *i* steps apart around the circle of demes, (*d*−*i*)*i/*(*M* +*X*) is the expected time, in units of 2*N* generations, until they are in the same deme [28, p. 170]. Thus, the mean coalescence time for a pair of lineages *i* steps apart is

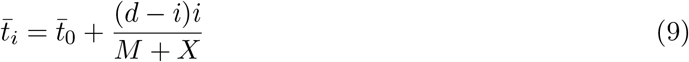

A pair in the same deme is not affected by extinction but can coalesce. Their expected coalescence time is

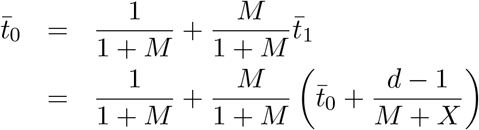

Solving for 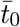 gives

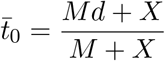

just as in the island model.

To calculate *F*_*ST*_, we need 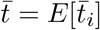, which depends on *E*[(*d*−*i*)*i*], where *i* is the distance between two genes drawn at random from the entire population. Suppose the first gene is from deme 0. The second is equally likely to come from demes 1, 2, …, *d*, where the demes are numbered clockwise around the circle, and deme *d* is the same as deme 0. If *i* represents the “clockwise distance” between demes, then the shortest distance from deme *i* to deme 0 is the minimum of *i* and *d* − *i*. Because we are averaging (*d* − *i*)*i*, it doesn’t matter whether we interpret *i* as the clockwise distance or the shortest distance: (*d* − *i*)*i* will be the same in either case. For convenience, I take *i* as the clockwise distance. The expectation of (*d* − *i*)*i* is

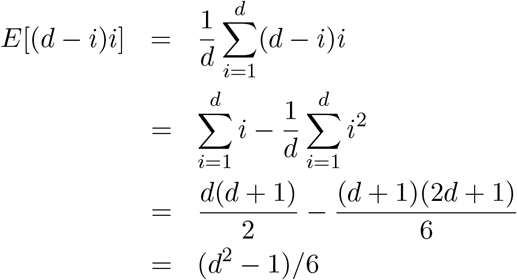

Substitute this for (*d* − *i*)*i* in Eqn. 9 to obtain

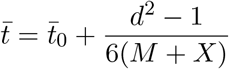

Now Eqn. 8 gives

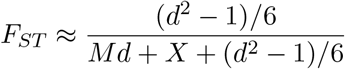

which is equivalent to Eqn. 6 above.

## References

[1] Franz Baumdicker et al. “Efficient ancestry and mutation simulation with Msprime 1.0”. Genetics 220.3 (Dec. 2021), iyab229. doi: 10.1093/genetics/iyab229.

[2] Mogens Bladt and Bo Friis Nielsen. Matrix-Exponential Distributions in Applied Probability. Springer, 2017. isbn: 978-1-4939-7049-0. doi: 10.1007/978-1-4939-7049-0.

[3] D. Carmelli and L.L Cavalli-Sforza. “Some models of population structure and evolution”. Theoretical Population Biology 9.3 (1976), pp. 329–359. doi: 10.1016/0040-5809(76)90052-6.

[4] Lounès Chikhi et al. “The IICR (inverse instantaneous coalescence rate) as a summary of genomic diversity: insights into demographic inference and model choice”. Heredity 120.1 (2018), pp. 13–24. doi: 10.1038/s41437-017-0005-6.

[5] D.R. Cox and H.D. Miller. The Theory of Stochastic Processes. London: Chapman and Hall, 1965.

[6] Richard R. Hudson. “Gene genealogies and the coalescent process”. In: Oxford Surveys in Evolutionary Biology. Ed. by Douglas Futuyma and Janis Antonovics. Vol. 7. Oxford: Oxford University Press, 1990, pp. 1–44.

[7] Guy S Jacobs et al. “Multiple deeply divergent Denisovan ancestries in Papuans”. Cell 177 (2019), pp. 1–12. doi: 10.1016/j.cell.2019.02.035.

[8] J. D. Kalbfleisch and R. L. Prentice. The Statistical Analysis of Failure Time Data. New York: Wiley, 1980. doi: 10.2307/3315078.

[9] Jerome Kelleher, Alison M Etheridge, and Gilean McVean. “Efficient coalescent simulation and genealogical analysis for large sample sizes”. PLoS Computational Biology 12.5 (May 2016), pp. 1–22. doi: 10.1371/journal.pcbi.1004842.

[10] Motoo Kimura and George H. Weiss. “The stepping stone model of population structure and the decrease of genetic correlation with distance”. Genetics 49 (1964), pp. 561–576.

[11] Valérie Laporte and Brian Charlesworth. “Effective population size and population subdivision in demographically structured populations”. Genetics 162.1 (2002), pp. 501–519. doi: 10.1093/genetics/162.1.501.

[12] Heng Li and Richard Durbin. “Inference of human population history from individual whole-genome sequences”. Nature 475.7357 (2011), pp. 493–496. doi: 10.1038/nature10231.

[13] Fabrizio Mafessoni et al. “A high-coverage Neandertal genome from Chagyrskaya Cave”. Proceedings of the National Academy of Sciences, USA 117.26 (2020), pp. 15132–15136. doi: 10.1073/pnas.2004944117.

[14] T. Maruyama and M. Kimura. “Genetic variability and effective population size when local extinction and recolonization of subpopulations are frequent”. Proceedings of the National Academy of Sciences, USA 77 (1980), pp. 6710–6714. doi: 10.1073/pnas.77.11.6710.

[15] Takeo Maruyama. “On the rate of decrease of heterozygosity in circular stepping stone models”. Theoretical Population Biology 1.1 (1970), pp. 101–119. doi: 10.1016/0040-5809(70)90044-4.

[16] Takeo Maruyama. Stochastic Problems in Population Genetics. New York: Springer-Verlag, 1977.

[17] O. Mazet et al. “On the importance of being structured: instantaneous coalescence rates and human evolution—lessons for ancestral population size inference?” Heredity 116.4 (2016), pp. 362–371. doi: 10.1038/hdy.2015.104.

[18] Olivier Mazet and Camille Noûs. “Population genetics: coalescence rate and demographic parameters inference”. Peer Community Journal 3.e53 (2023). doi: 10.24072/pcjournal.285.

[19] Masatoshi Nei and Naoyuki Takahata. “Effective population size, genetic diversity, and coalescence time in subdivided populations”. Journal of Molecular Evolution 37 (1993), pp. 240–244. doi: 10.1007/BF00175500.

[20] Willy Rodríguez et al. “The IICR and the non-stationary structured coalescent: towards demographic inference with arbitrary changes in population structure”. Heredity 121.6 (2018), pp. 663–678. doi: 10.1038/s41437-018-0148-0.

[21] Alan R. Rogers. “Legofit: estimating population history from genetic data”. BMC Bioinformatics 20 (2019), p. 526. doi: 10.1186/s12859-019-3154-1.

[22] Alan R. Rogers. “An efficient algorithm for estimating population history from genetic data”. Peer Community Journal 2 (2022), e32. doi: 10.24072/pcjournal.132.

[23] François Rousset. Genetic Structure and Selection in Subdivided Populations. Princeton University Press, 2004. isbn: 0691088179. doi: 10.1515/9781400847242.

[24] Stephan Schiffels and Richard Durbin. “Inferring human population size and separation history from multiple genome sequences”. Nature Genetics 46.8 (Aug. 2014), pp. 919–925. doi: 10.1038/ng.3015.

[25] Jason P. Sexton, Sandra B. Hangartner, and Ary A. Hoffmann. “Genetic isolation by environment or distance: which pattern of gene flow is most common?” Evolution 68.1 (Jan. 2014), pp. 1–15. doi: 10.1111/evo.12258.

[26] Montgomery Slatkin. “Gene flow and genetic drift in a species subject to frequent local extinctions”. Theoretical Population Biology 12 (1977), pp. 253–262.

[27] Montgomery Slatkin. “The average number of sites separating DNA sequences drawn from a subdivided population”. Theoretical Population Biology 32.1 (1987), pp. 42–49. doi: 10.1016/0040-5809(87)90038-4.

[28] Montgomery Slatkin. “Inbreeding coefficients and coalescence times”. Genetical Research, Cambridge 58 (1991), pp. 167–175.

[29] Curtis Strobeck. “Average number of nucleotide differences in a sample from a single subpopulation: a test for population subdivision”. Genetics 117.1 (1987), pp. 149–153. doi: 10.1093/genetics/117.1.149.

[30] Yuri Sukhov and Mark Kelbert. Probability and Statistice by Example: II: Markov Chains: A Primer In Random Processes and their Applications. Cambridge: Cambridge University Press, 2008.

[31] Simon Tavaré. “Line-of-descent and genealogical processes, and their applications in population genetics models”. Theoretical Population Biology 26 (1984), pp. 119–164. doi: 10.1016/0040-5809(84)90027-3.

[32] Jonathan Terhorst, John A Kamm, and Yun S Song. “Robust and scalable inference of population history from hundreds of unphased whole genomes”. Nature Genetics 49.2 (2017), pp. 303–309. doi: 10.1038/ng.3748.

[33] John Wakeley. “Nonequilibrium migration in human history”. Genetics 153.4 (1999), pp. 1863–1871. doi: 10.1093/genetics/153.4.1863.

[34] M. C. Whitlock and Nicholas H. Barton. “The effective size of a subdivided population”. Genetics 146 (1997), pp. 427–441. doi: 10.1093/genetics/146.1.427.

[35] M. C. Whitlock and D. E. McCauley. “Some population genetic consequences of colony formation and extinction: genetic correlations within founding groups”. Evolution 44.7 (1990), pp. 1717–1725. doi: 10.1111/j.1558-5646.1990.tb05243.x.

[36] Hilde M. Wilkinson-Herbots. “Genealogy and subpopulation differentiation under various models of population structure”. Journal of Mathematical Biology 37.6 (1998), pp. 535–585. doi: 10.1007/s002850050140.

[37] Sewall Wright. “Breeding structure of populations in relation to speciation”. American Naturalist 74.752 (1940), pp. 232–248. doi: 10.1086/280891.

[38] Sewall Wright. “Isolation by distance”. Genetics 28.2 (1943), pp. 114–138. doi: 10.1093/genetics/28.2.114.

